# Near 100% genome editing efficiency with a one-step all RNA prime editing system in a human reporter iPSC line

**DOI:** 10.1101/2024.12.20.629762

**Authors:** Kyle Wettschurack, William C. Skarnes, Enrica Pellegrino, Morgan Robinson, Yang Yang

**Author notes:** These authors contributed equally to this work (Kyle Wettschurack, William C. Skarnes).

## Abstract

CRISPR/Cas9 nucleases offer powerful tools for genome editing but can cause unintended mutations at the target site due to the creation of double-stranded DNA breaks. Prime editing (PE) is arguably a safer technology as it relies on the creation of single-strand DNA nicks and avoids possible indels. Despite its precision, current PE systems are limited by relatively low editing efficiency in non-immortalized cells. Here, we optimized prime editing in human induced pluripotent stem cells (hiPSCs) using a fluorescence-based assay to achieve near-complete editing of a BFP transgene. Using an all-RNA approach, precise prime editing of a single nucleotide variant was observed in >95% of cells with minimal unintended edits at the target site.

## Introduction

Genome editing technologies have advanced remarkably in recent years, ushering in an era of unprecedented precision and versatility in genetic research. Among these, clustered regularly interspaced short palindromic repeats (CRISPR)-Cas systems have been widely adopted for their ability to create precise DNA modifications using programmable guide RNAs^1,2^. While the use of a traditional Cas9 nuclease-based approaches, combined with a homology-directed repair (HDR) template, can achieve high editing efficiencies^3,4^, their reliance on inducing double-stranded DNA breaks (DSBs) poses significant limitations, including the risk of unwanted indels, off-target effects, and genomic rearrangements at the target site. To address these limitations, CRISPR Prime Editing (PE) was developed as a safer and more precise alternative^5^. Prime editing employs a Cas9 nickase fused to a reverse transcriptase enzyme, introducing single-strand DNA nicks instead of double-strand breaks. These nicks are repaired using the sequence encoded by a prime editing guide RNA (pegRNA), enabling precise insertions, deletions, or nucleotide substitutions. Unlike CRISPR base editing, which can potentially result in unintended “bystander edits^6^”, prime editing minimizes such off-target effects^5,7^. Despite these advantages, current PE systems often suffer from relatively low editing efficiencies. Recent progress in prime editing has incorporated strategies such as secondary nicking guide RNAs (ngRNAs)^5^, mismatch repair inhibitors^7^, and engineered prime editing guide RNAs^8^ (epegRNAs), which have shown promise in improving outcomes.

Human induced pluripotent stem cells (hiPSCs) have emerged as a preferred cell type for modeling human diseases^9^. Derived from reprogrammed adult somatic cells, such as fibroblasts or blood cells, hiPSCs are pluripotent cells, enabling their differentiation into a wide array of human cell types^10^. This versatility makes hiPSCs invaluable for disease modeling, genetic screening, and therapeutic applications. Achieving high editing efficiency in hiPSCs is critical for creating disease models, generating isogenic controls from patient lines, and performing robust genetic screens. However, prime editing efficiencies in hiPSCs remain suboptimal. Previous studies have reported efficiencies of approximately 25%, with iterative re-editing increasing efficiencies to around 50% but at the cost of significant indel formation^11^. More recently, single-step transfection methods have demonstrated editing efficiencies of ∼50% in hiPSCs, although substantial variability persists^12^. Here, we present our optimization of prime editing in a hiPSC line containing a fluorescent reporter, designed to monitor the efficiency of single nucleotide mutations. Using nucleofection to deliver RNA-encoded prime editing reagents, we achieved nearly 100% editing efficiency with minimal unintended edits. These results underscore the potential to achieve near-perfect editing efficiencies with minimal off target edits.

## Results

### Improving editing efficiency in a BFP-GFP assay with a plasmid-based prime editor

To quickly and efficiently optimize prime editing in hiPSCs, we used a fluorescence-based flow cytometry assay in a hiPSC line that contains a constitutively expressed blue fluorescent protein (BFP) inserted in the AAVS1 site^3^. BFP can be converted to green fluorescent protein (GFP) by introducing a single C>T nucleotide change, converting histidine to tyrosine at position 67 (H67Y)^4^ (**Fig. 1A and Supp. Fig. 1A-B**). AAVS1-BFP hiPSCs stably express BFP in 99% of cells over several passages while less than 1% show no fluorescence (**Supp. Fig. 1C**). We started by nucleofecting AAVS1-BFP cells with PEmax plasmid^7^ (addgene #174820) encoding the prime editor and MutL homolog dominant negative (MLH1dn) protein and a synthetic epegRNA designed to create the H67Y amino acid change (**Supp. Table 1**). Six days post-nucleofection, we observed a low level of prime editing where only 4% of cells express GFP (**Fig. 1B**). We observed two distinct populations of GFP-positive cells, GFP (high) and GFP/BFP double-positive cells, not seen in previous experiments using Cas9 nuclease and a repair oligo to introduce the H67Y amino acid change^3,4^. Cell sorting and molecular characterization of these two GFP populations revealed the presence of two copies of the BFP transgene in one allele of AAVS1 (**Supp. Fig. 1**). Sanger sequencing and PCR analysis confirmed that both copies of BFP are edited in GFP(high) sorted cells, whereas only one copy of GFP is modified in GFP(low)/BFP double positive cells. Thus, fortuitously, the insertion of a tandem duplication of the BFP donor plasmid in AAVS1-BFP cells allows us to monitor prime editing of one versus two copies of the target sequence.

**Figure 1.**
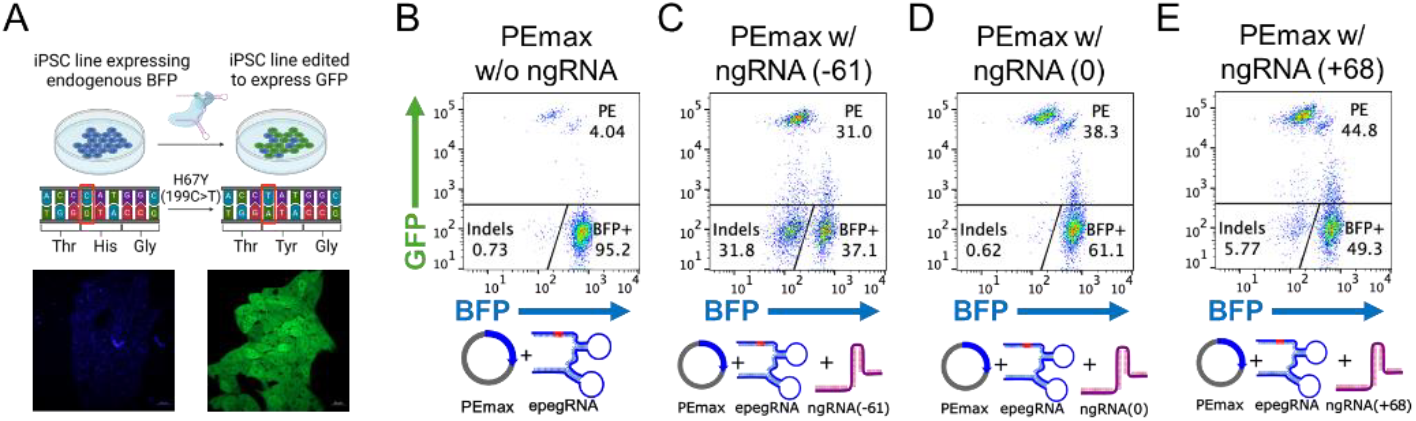
BFP-GFP editing in iPSCs using plasmid: **A)** Schematic of BFP-> GFP editing (top). Images of hiPSCs expressing BFP and GFP (bottom). **B)** BFP->GFP FACS results using PEmax plasmid without a ngRNA. **C)** FACS plot using PEmax plasmid with the addition of a -61 ngRNA (the ngRNA nicks 61 bases upstream of the nick created by the epegRNA). **D)** FACS plot using PEmax plasmid with the addition of a 0 ngRNA (the ngRNA nicks at the same location of the nick created by the epegRNA). **E)** FACS plot using PEmax plasmid with the addition of a +68 ngRNA (the ngRNA nicks 68 bases downstream of the nick created by the epegRNA).

Nicking of the opposite strand is known to increase the efficiency of prime editing^5,7^. We designed three different ngRNAs to introduce nicks at positions -61, 0, and +68 relative to the nick created by the epegRNA spacer (**Supp. Table 1**). Nicking at -61 yielded a 10-fold increase (31.0%) in prime editing efficiency, accompanied by a high loss of BFP signal (31.8%) presumably caused by indel formation at double-strand breaks (**Fig. 1C**). The use of ngRNA(0), which overlaps the epegRNA, showed prime editing in 38% of cells without a measurable incidence of indels (**Fig. 1D**), whereas nicking by ngRNA(+68) produced the highest prime editing in 45% of cells with a modest (6%) incidence of indel formation (**Fig. 1E**). These differences in indel formation using guides at different positions are consistent with previous results^7^ and dual nicking studies^13,14^. Both double-strand breaks and indel formation are common outcomes when using Cas9 nickase with guide RNAs (gRNAs) arranged in a tail-to-tail orientation ^13,14^.

### Increasing editing efficiency in a BFP-GFP assay with an all-RNA approach

Recent work has demonstrated improved prime editing efficiencies using RNA delivery of prime editing components^11,12^. We next performed the BPF-GFP assay by co-nucleofecting increasing amounts of PEmax and dnMLH1 mRNA, along with epegRNA. Significant cell death was observed in nucleofections containing more than 16 ug of PEmax (data not shown). When keeping PEmax mRNA constant, we did not observe any improvement in prime editing efficiency by increasing the amount of dnMLH1 mRNA or epegRNA (data not shown). The optimal amount of each component for a 20 microliter nucleofection was determined empirically to be 16 ug PEmax mRNA, 6 ug MLH1dn mRNA (a 1:1 molar ratio with PEmax), and 8 ug epegRNA. In the absence of a nicking guide, 36% of cells express GFP, with a predominance of the GFP(low)/BFP double-positive cell population (**Fig. 2A**), representing a ∼10-fold increase in prime editing efficiency compared to plasmid delivery (**Fig. 1B**).

**Figure 2.**
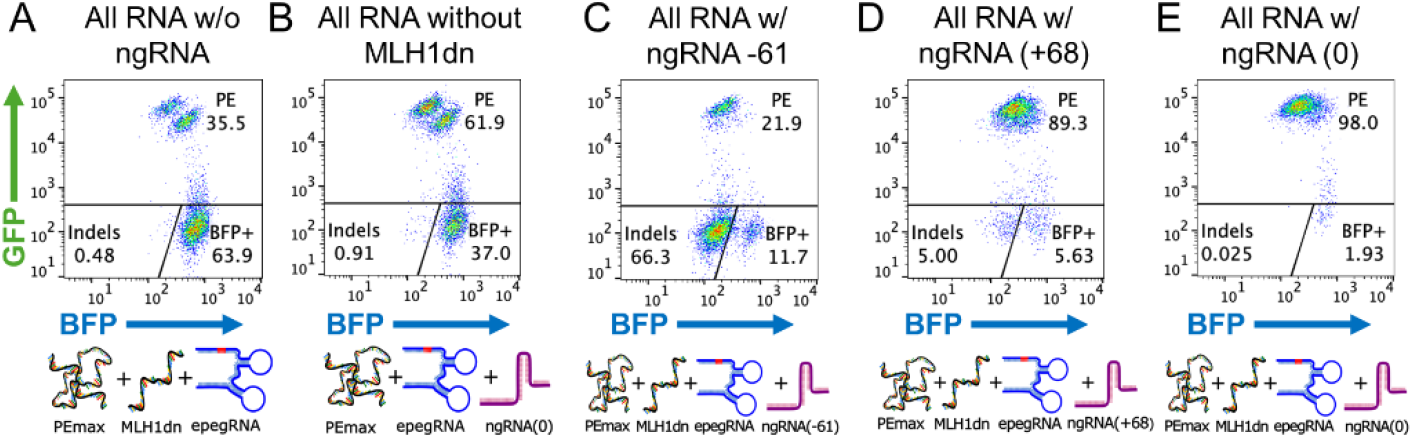
BFP-GFP editing in hiPSCs using an all-RNA approach: **A)** FACS plot of prime editing delivering PEmax and MLH1dn as *in vitro* transcribed mRNA, this reaction did not contain a ngRNA. **B)** FACS result of prime editing using an all-RNA approach without MLH1dn and using ngRNA(0). **C)** FACS result of prime editing using an all-RNA approach and adding the ngRNA(−61). **D)** FACS result of prime editing using an all-RNA approach and adding the ngRNA(+68). **E)** FACS result of prime editing using an all-RNA approach and adding the ngRNA(0).

We then measured the effect of nicking guide RNAs on prime editing efficiency. The addition of ngRNA(0) while removing MLH1dn resulted in an editing efficiency of 62% (**Fig. 2B**) Addition of ngRNA(−61) resulted in a dramatic loss of BFP fluorescence in 66% of cells (**Fig. 2C**), again highlighting the importance of avoiding ngRNAs in a tail-to-tail orientation. The addition of ngRNA(+68) significantly improved prime editing to 89%, with modest indel formation in 5% of cells (**Fig. 2D**), whereas ngRNA(0) resulted in prime editing in a remarkable 98% of cells (**Fig. 2E**) with no detectable indels.

## Discussion

In this study, we optimized prime editing in a human reporter hiPSC line using a BFP-to-GFP conversion assay, achieving near 100% editing efficiency in a single nucleofection with no detectable on-target indels. Our protocol is highly reproducible, as evidenced by >95% editing efficiency in two independent laboratories, performed by different operators using commercially-sourced reagents. To gain optimal prime editing efficiency in our RNA only experiments, we varied the amount of each reagent (data now shown). In a 20 microliter nucleofection, we increased the amount of PEmax mRNA from 4 ug to 32 ug, keeping the molar ratio between PEmax and dnMLH1 mRNA constant at 1:1, and found that the cell viability decreases significantly at greater than 16 ug of the editor. We also saw this effect when there was greater than 2 ug of the PEmax plasmid. Increasing the molar ratio of PEmax and MLH1dn to 1:3 had no significant effect on editing efficiency (data not shown). Increasing the amount of epegRNA and ngRNA (> 8 and 5 ug respectively) also had little effect on efficiency. We compared capped, synthetic mRNAs with and without 5-methoxyuridine (5moU) modification and found that unmodified mRNA performed slightly better in our hands (data not shown). Finally, we examined whether or not adding a silent mutation to both the epegRNA and overlapping ngRNA(0) increased editing efficiency^7^, but no benefit was seen with the silent mutation (data not shown).

Our approach leverages PEmax, an improved version of the PE2 editor. PEmax incorporates two mutations in the SpCas9 sequence, a codon-optimized reverse transcriptase domain, and two additional nuclear localization sequences (NLS), enhancing its editing ability^7^. Recent iterations, such as PE6^15^ and PE7^16^, offer unique modifications, but their compatibility with our optimized method remains to be investigated. Beyond the editor itself, our study underscores the critical role of other prime editing components. For example, blocking mismatch repair through the expression of MLH1dn significantly improved editing efficiency. By far, the introduction of ngRNA had the largest effects on prime editing efficiency and on countervailing indel formation. The placement of a nick on the strand opposite of the epegRNA proved crucial: ngRNA overlapping the mutation site yielded the highest editing efficiency with no detectable indels, as measured by flow cytometry. Our data indicates that placing a ngRNA in the correct position results in no increase in indels compared to a reaction lacking a ngRNA. Introducing a nick downstream of the epegRNA (head-to-head orientation) also achieved high-efficiency prime editing with a relatively low percentage of indels (5%). In contrast, a nick upstream of the epegRNA (tail-to-tail configuration) should be avoided due to their tendency to create double-strand breaks that are then readily converted to indels^13,14^.

Near 100% editing efficiency implies near 100% efficiency of delivery. Here, we utilized nucleofection, a delivery method ideal for hiPSCs but less well-suited to other cell types, such as neurons. Efficient delivery of editing reagents remains a major challenge for the application of genome editing technology to additional cell types both *in vitro* and *in vivo*. It is not known if high efficiency editing can be achieved in clinically relevant targets. For example, does prime editing efficiency decrease in post-mitotic cells like neurons, even with optimal delivery systems? Could chromatin accessibility and other factors influence editing outcomes? We are currently conducting experiments to address these questions and remain optimistic that improved methods can achieve high editing efficiency in disease-relevant targets across a broader range of cell types.

**Supplementary Fig 1.**
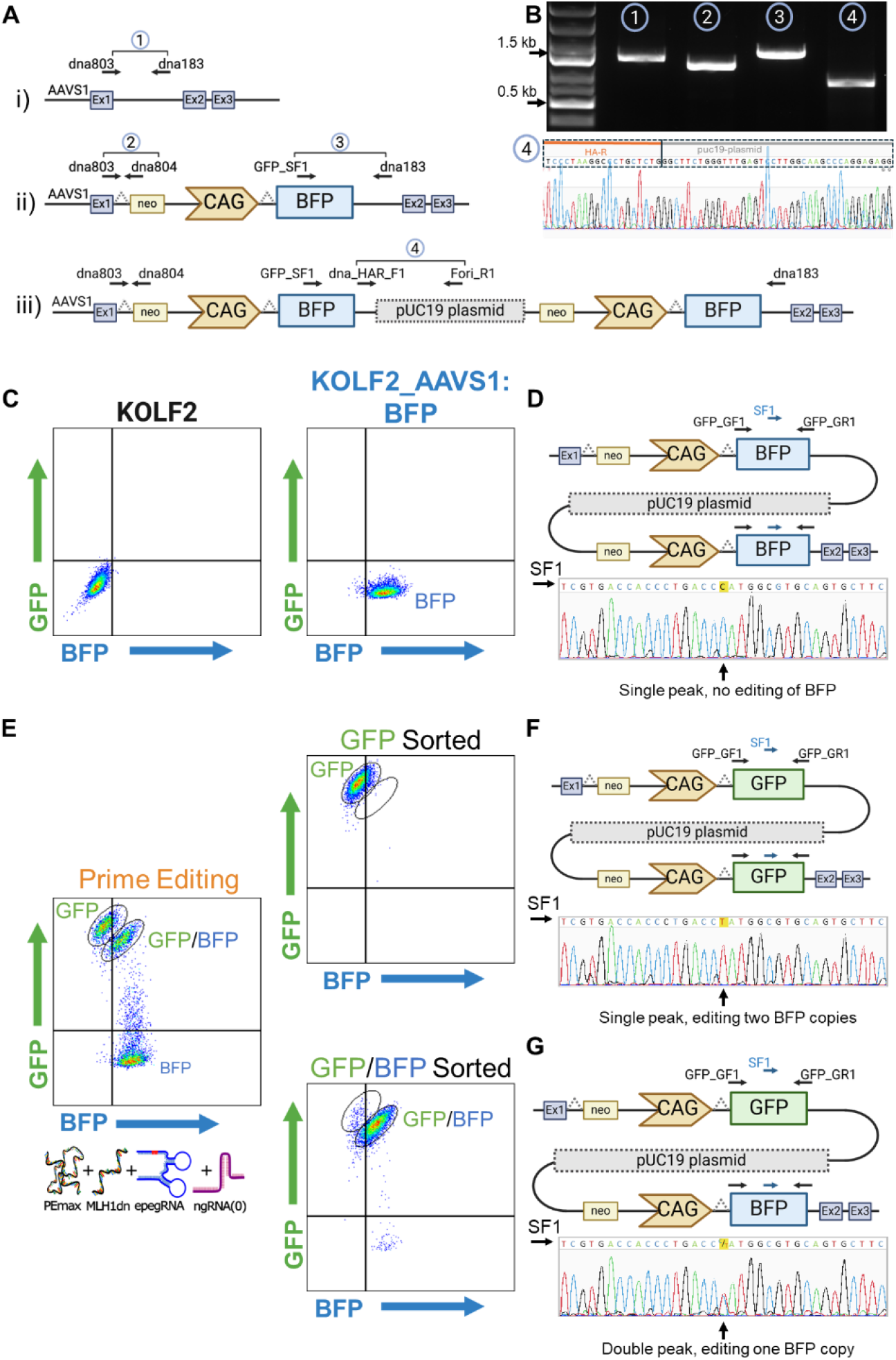
Characterization of AAVS1:BFP hiPSCs and prime edited pools: **A)** (i) Schematic of the AAVS1 locus and (ii) the neo-CAG-BFP sequence in the AAVS1 locus of the Kolf2.1 cells. PCR genotyping primers are indicated with black arrows and PCR primer pairs and amplicon sizes numbered (1) – (4). Gray boxes indicate PPP1R12C exons (Ex1, Ex2 and Ex3). (iii) Hypothetical representation of a tandem integration of the neo-CAG-BFP donor plasmid containing the pUC19 plasmid backbone. **B)** Top. PCR genotyping of the AAVS1:BFP hiPSC clone confirming the presence of a wild-type (non-targeted) allele (reaction 1), a targeted allele (reactions #2 and 3) and the presence of the pUC plasmid backbone (reaction #4). Bottom. Sanger sequencing of unedited cells KOLF2_AAVS1:BFP cells shows pUC19 plasmid integration. **C)** Flow cytometry of KOLF2.1 control cells (no BFP) and KOLF2.1_AAVS1:BFP cells. **D)** Schematic representation of donor plasmid integration containing two copies of the neo-CAG-BFP transgene. Sanger sequencing of unedited cells shows BFP sequence only. **E)** Flow cytometry of AAVS1:BFP cells after prime editing with PEmax, MLH1dn, epegRNA and ngRNA(0). GFP(high) and GFP/BFP populations are circled before and after cell sorting. **F)** Schematic representation of donor plasmid integration containing two copies of GFP following prime editing. Sanger sequencing of GFP(high) cells shows GFP sequence only. **G)** Schematic representation of donor plasmid integration containing one copy of BFP and one copy of GFP. Sanger sequencing of GFP/BFP cells shows both the GFP and BFP sequence.

**Supplementary table 1:**
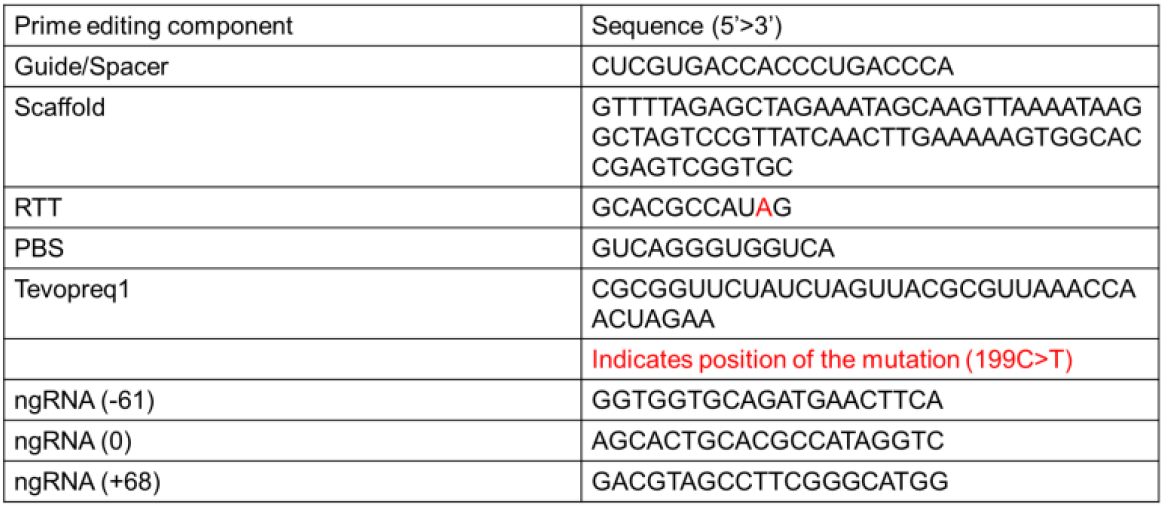
Sequence of the regions of the epegRNA and the ngRNAs used in this study.

## Methods

### epegRNA design and *in-vitro* Transcribed Editing Material

epegRNA’s were designed manually based on design guidelines described in the literature^8,17,18^. The guide section is also a 20-base pair sequence adjacent to an NGG PAM site. The scaffold sequence was consistent with literature^5^. The RTT region used for these experiments was 11 nucleotides while the PBS was 13 nucleotides. The tevopreq1 sequence was appended to the 3’ end of the epegRNA. The epegRNA was custom-synthesized by Integrated DNA Technologies (IDT). The first 3 bases and the last two bases of the epegRNA were Phosphorothioated and 2’ O-methyl modified. ngRNA’s were purchased from Synthego as ‘standard gRNA’ in tubes. The gRNA contained the standard modifications (2’-O-Methyl at 3 first and last bases, 3’ phosphorothioate bonds between first 3 and last 2 bases). PEmax and MLH1dn were custom synthesized from Trilink Biotechnologies based on the sequences of PEmax and MLH1dn in addgene plasmid #174820. Both PEmax and MLH1dn sequences used unmodified mRNA with wild-type bases and CleanCap® technology.

### hiPSC culture

hiPSCs were cultured in StemFlex (Gibco A3349401) in a 6-well plate (Corning 3516) coated with Synthemax (Sigma-Aldrich CLS3535). When the cells reached 70%-80% confluency the cells were passaged by first washing with 1 mL of PBS, adding 2 mL of ReLeSR (STEMCELL Technologies 100-0483) to the well, and incubating at 37°C and 5% CO_2_ for 5 minutes. The ReLeSR was removed and 3 mL of room-temperature StemFlex was added to the well. The cells were then removed from the bottom of the plate using a cell lifter (Corning 3008). Synthamax was removed and 3 mL of StemFlex was added to one well of a 6-well plate. The cells were then passaged at a dilution of 1:15 into the new 6-well plate. StemFlex media was changed every 2 days for optimal cell health.

### Ethanol precipitation of editing materials

For each nucleofection reaction using plasmid 2 ug of plasmid was used. For the reactions using in-vitro transcribed mRNA 16 ug of PEmax RNA and 6 ug of MLH1dn were used. These items were resuspended in P3 nucleofection buffer after ethanol precipitation. To perform this precipitation the editing material was mixed with 500 uL of 100% EtOH, 180 uL of H_2_O, and 4 uL of 5M NaCl. After all components were added, the tube was mixed by inverting several times. This solution was kept at -20°C for 1 hour and then spun at 15,000 RPM for 30 minutes at 4°C. The supernatant was then decanted and the pellet was gently washed three times with 1 mL of 70% EtOH. After this, the pellet was left to air dry for 5-10 minutes until the ethanol was evaporated. The pellet was then resuspended in 18 uL P3 buffer (Lonza V4XP-3032) and left at 4°C overnight.

### Preparation of the P3 nucleofection solution

Before nucleofection, we added 4 uL of supplement 1 (Lonza V4XP-3032) to the P3 solution containing the resuspended editing material. Then we added 8 ug of epegRNA and 5.2 ug of ngRNA to the reaction (the same amount of epegRNA and ngRNA was used in both plasmid and RNA-based reactions).

### Nucleofection

Nucleofections were carried out similarly to our protocol for hiPSCs nucleofection^*3*^. hiPSCs were cultured in StemFlex for at least one passage after thawing. Two hours before beginning the experiment, we performed a full media change with StemFlex. After two hours, we washed the cells once with 2 mL of PBS. Then we dissociated the cells using Accutase (STEMCELL Technologies 07920) and centrifuged them at 300xg for 5 minutes in 3 mL of StemFlex + RevitaCell supplement (ThermoFisher A2644501). Cells were then resuspended in 1 mL of StemFlex media + RevitaCell. 2×10^5^ cells from the suspension were counted using a hemocytometer. These cells were then added to a v-bottom plate and spun at 300xg for 5 minutes. After this the media was aspirated and the pellet was resuspended in 20 uL of P3 solution containing the PEmax plasmid or in-vitro transcribed mRNA, epegRNA, and ngRNA if applicable. The P3/cell solution was placed in a 16-well strip and nucleofected using an Amaxa 4D nucleofector (Lonza 4D-Nucleofector X Unit AAF-1003X) with CA137 settings. After nucleofection, cells were replated on a vitronectin coated 24 well plate in StemFlex and CloneR2 (STEMCELL Technologies 100-0691).

### FACS Sorting

StemFlex was changed 48 hours after nucleofection to remove CloneR2, and cells were grown in StemFlex for 4-6 days before sorting, with full media changes every other day and passaging into a 6-well plate when cells reached 70-80% confluency. For FACS sorting, cells were passaged with Accutase and resuspended in StemFlex plus CloneR2, then sorted using a FACSymphony S6 CT using a nozzle size of 100 um with a speed of 1.0. The results were analyzed using FlowJo software.

